# Exploring Behavioral and Neural Dynamics of Social Interactions during the Juvenile Stage in a Mouse Model of Fragile X Syndrome

**DOI:** 10.1101/2024.03.03.583140

**Authors:** Moon Young Bae, Bosong Wang, Asim Ahmed, Abdullah Abdullah, Raffay Ilyas, Veronica Rasheva, Kartikeya Murari, Ning Cheng

**Author notes:** Equal contribution (Moon Young Bae, Bosong Wang).

## Abstract

**Introduction:** Fragile X Syndrome (FXS), caused by *Fmr1* mutations, is linked to cognitive and behavioral differences, including altered social interactions. Most mouse studies focus on adults, despite human research showing critical developmental changes in childhood and adolescence. We examined social behavior in juvenile male and female *Fmr1* knockout (KO) mice as well as heterozygous (HET) females. We further assessed cortical activity in KO females to better understand early phenotypes.

**Methods:** Juvenile mice of both sexes and genotypes were paired in same-sex, novel dyads for 10-minute interactions. Key social behaviors such as head, anogenital, and body sniffing, and physical touch, as well as distance travelled, were analyzed with a marker-less tracking software. Frontal-parietal EEG recordings were collected from wild-type (WT) and KO females in home cage and social contexts to analyze power spectra across frequency bands.

**Results:** HET and KO females engaged in more frequent but shorter interaction events compared to WT females, with HET females showing the highest counts. Males displayed similar trends when comparing KO and WT. Males engaged in overall higher interaction events than females. EEG analyses revealed altered oscillatory activity in KO females compared to WT females, especially within theta, alpha, and beta bands, most prominently during the early interaction phase. Locomotor activity correlated weakly with head/anogenital sniffing but more strongly with body sniffing and touch.

**Discussion:** These findings suggest that *Fmr1*-related differences in juvenile social behavior are sex-dependent and associated with cortical oscillatory changes. Characterizing these early phenotypes in both sexes allows us to further understand FXS development and informs potential routes for early intervention.

**Key Points:**

- Fragile X Syndrome (FXS), the leading inherited cause of autism, is associated with disruptions in social behavior.
- While social phenotypes are relatively well described in adult mouse models of FXS, juvenile manifestations remain poorly understood.
- Social behavior was assessed in juvenile male and female *Fmr1* knockout (KO), heterozygous (HET, female only), and wildtype (WT) mice, and frontal-parietal EEG recordings were collected from WT and KO females.
- HET and KO females exhibited more frequent but shorter social interactions than WT females, with HET showing the greatest number of events. Males showed similar patterns when comparing KO and WT. Males engaged in higher overall interaction events than females. EEG recordings revealed altered oscillatory activity in KO compared to WT females, most pronounced during the early phase of social encounter.
- These findings reveal sex- and genotype-dependent differences in juvenile social behavior and cortical activity, highlighting the importance of studying juvenile development in FXS.

## 1. Introduction

FXS is the leading monogenic cause of autism spectrum disorder (ASD), with an estimated prevalence of 1 in 4,000 males and 1 in 8,000 females (Hunter *et al*., 1993). It occurs due to an expansion in the number of CGG trinucleotide repeats within the 5’-untranslated region of the Fragile X Messenger Ribonucleoprotein 1 (*Fmr1*) gene (Hunter *et al*., 1993). This leads to the subsequent loss of the Fragile X Messenger Ribonucleoprotein (FMRP), which causes widespread dysregulation of synaptic function and plasticity, particularly in regions that process external cues such as the prefrontal cortex, amygdala, and hippocampus (Cregenzán-Royo *et al*., 2022; Olmos-Serrano & Corbin, 2011; Telias, 2019; Sidorov *et al*., 2013; Tian *et al*., 2017). FXS is associated with cognitive and behavioral differences, including hyperactivity, heightened anxiety-like responses in social settings, and decreased brain activation in areas such as the amygdala that mediate social cognition (Selmeczy *et al*., 2011; Shelton *et al*., 2017).

Social behavior encompasses any form of communication or interaction between two or more individuals and is crucial for adapting to a new environment (Jabarin *et al*., 2022). Social interactions are closely tied to an individual’s emotional state, presenting challenges for research into the underlying mechanisms (Jabarin *et al*., 2022). Difficulties with social interactions during development are common in FXS (Cregenzán-Royo *et al*., 2022). Cregenzan-Royo et al. (2022) reported that about 21.5% and 17.5% of boys and girls with FXS, respectively, between 6 and 17 years of age, displayed social withdrawal, while another study by Hessl *et al*. (2001) showed that 40% of girls and 41.8% of boys (aged 6-16 years) with FXS had social problems in the clinical range. Poor socialization skills have been associated with adverse outcomes, including symptoms of anxiety and/or depression, reduced adaptive skills, and decreased quality of life (Lightbody *et al*., 2022). Despite showing genuine interest in social engagement, many experience anxiety and exhibit avoidance behaviors in unfamiliar settings (Roberts *et al*., 2007). For instance, males with FXS commonly avoid eye contact and physically orient their bodies away from others (Kau *et al*., 2004). High occurrence of intellectual disability in FXS in attention or working memory further contribute to difficulties in social communication (Kau *et al*., 2004).

Research indicates that behavior challenges in FXS emerge during infancy, intensify throughout childhood, and stabilize during adolescence and early adulthood (Roberts *et al*., 2019). Approximately 81% of males with FXS displayed social avoidance from ages 4 months to 25 years (Roberts *et al*., 2019). Findings from a study by Black *et al*. suggest that both behavioral and physiological signs of social anxiety were evident at 12 months of age in infants with FXS (Black *et al*., 2021). Young boys with FXS aged 36 to 95 months also show slower development of socially adaptive behaviors (Bailey *et al*., 2000). Compared to typically developing children, those with FXS display heightened anxiety, gaze aversion, and social withdrawal during challenging social tasks (Klusek *et al*., 2020; Kau *et al*., 2004). Social difficulties, anxiety, and adaptive behavior problems remain a prevalent concern for individuals aged 3 to 30 years (Côté *et al*., 2021). While research on adolescent humans with FXS is extensive, animal studies in FXS during early development are limited (Gauducheau *et al*., 2017). Early investigation is crucial for understanding the developmental trajectory of social and cognitive symptoms and identifying potential therapeutic interventions. Given that FXS represents a major genetic risk factor for ASD, such knowledge could help with the implementation of early therapeutic interventions for children with ASD (Oddi *et al*., 2015).

Several studies have utilized the *Fmr1* knockout (KO) mouse model of FXS to investigate social behaviors and neural activity (Saré *et al*., 2019). This is because they exhibit certain similar traits in the human syndrome, such as hyperactivity, intellectual and cognitive differences, as well as reduced preference for novelty in social settings (Kazdoba *et al*., 2014). Previous studies by Spencer *et al*. (2005) showed that social interactions in male *Fmr1* KO mice at 3-4 months of age were experience-dependent, with greater social engagement observed among familiar mice. Social dominance tests found that 3-4-month-old male *Fmr1* KO mice retreated more frequently than WT mice (Spencer *et al*., 2005).

Recent research has explored electroencephalogram (EEG) as a promising biomarker for neurodevelopmental disorders (Kenny *et al*., 2022). EEG provides real-time feedback on cortical activity across regions linked to behavioral phenotypes, offering insights into the neural circuit dysfunctions in FXS (Kenny *et al*., 2022). EEG abnormalities, such as increased baseline gamma and theta power, along with reduced alpha power, have been observed in individuals with FXS and are believed to reflect underlying neural circuit dysfunctions (Wang *et al*., 2017). Developmentally, EEG patterns change, with children and adolescents showing exaggerated beta to gamma (∼ 25-50 Hz) power compared to typically developing peers (Wilkinson & Nelson, 2021). Preclinical studies in rodents have further linked EEG abnormalities to behavioral phenotypes in FXS. Findings include increased gamma power and behavioral traits such as hyperactivity and sensory hypersensitivity, paralleling human FXS features (Kozono *et al*., 2020; Wen et al., 2019).

FXS, being an X-linked disorder, exhibits more pronounced cognitive impairments in males (Cregenzán-Royo *et al*., 2022). As a result, most research has largely focused on males. However, there is a growing recognition of sex distinctions in social interactions observed in FXS, and the necessity to closely examine social behaviors and their manifestations in the female population. FXS phenotypes in females often present with learning disabilities, behavioral challenges, shyness, anxiety, and depression (Hunter *et al*., 1993). Regarding social skills, girls with FXS between the ages of 7 and 18 years displayed significantly lower rates in tasks, like identifying the causes and consequences of social problems (Russo-Ponsaran *et al*., 2014). Longitudinal research by Lightbody *et al*. (2022) found that depressive symptoms and social avoidance worsened with age in female patients, directly impacting social communication, reciprocal social behavior, and adaptive functioning. Interestingly, a previous study with female heterozygous *Fmr1* KO (HET) mice observed a hyper-social phenotype at 7-8 weeks of age, characterized by increased affiliative behaviors such as sniffing and physical contact (Petroni *et al*., 2022).

In terms of EEG, one human study found consistent alterations in theta and low beta power across sexes, while relative power differed by sex in the alpha, upper beta, gamma, and epsilon frequency bands (Smith *et al*., 2021). However, EEG studies in relation to social interaction particularly in females remain largely unexplored. The first aim of this study was to analyze the effects of genotype and sex on social behavior parameters in juvenile female and male *Fmr1* KO compared with HET females and wildtype (WT) control. To do so, we utilized a free-form social interaction test, and quantified interactions over time across multiple key facets of social behavior using a recently developed marker-less tracking algorithm (Le *et al*., 2024). To model the challenges individuals with FXS face in novel situations, mice were exposed to an unfamiliar environment to capture responses to novelty rather than habituation. We also aimed to characterize EEG spectral changes during the free social interactions between *Fmr1* KO and WT female groups. By integrating behavioral and neural data, we sought to better understand how neural oscillatory activity corresponds to social behavior changes in FXS.

## 2. Methods

### 2.1 Animals

Breeder wildtype (C57BL/6J, Jax stock No: 000664) and *Fmr1* KO (B6.129P2-*Fmr1^tm1Cgr^*/J, Jax stock No: 003025) mice were obtained from the Jackson Laboratory (ME, USA) and maintained at the mouse facility of the Cumming School of Medicine, University of Calgary. Behavioral testing was performed on male and female *Fmr1* KO and WT mice, bred in our animal facility at the University of Calgary. All KO mice of the current study were generated by mating *Fmr1* KO *(Fmr1 -/-)* homozygous females and *Fmr1* KO *(Fmr1 -/y)* males. For the heterozygous KO females in this study, mating involved wild-type (*Fmr1* +/+) C57BL/6 females with *Fmr1* KO (*Fmr1* -/y) males. Mice were group housed (up to five per cage) with their same-sex littermates. The cages were in a room with a controlled 12-hr light-dark cycle (lights on at 7:00am) with access to food and water ad libitum. Mouse pups were weaned around 20 days of age. All mice were fed standard mouse chow. Behavioral testing was performed between 09:00h and 19:00h. All behavioral testing procedures were performed in accordance with the guidelines in the Canadian Council for Animal Care and was approved by the Health Sciences Animal Care Committee of the University of Calgary.

### 2.2 Social Interaction Behavioral Test

For this experiment, 9 pairs of male WT mice, 11 pairs of male *Fmr1* KO mice, 7 pairs of female WT mice, 6 pairs of female *Fmr1* HET mice, and 6 pairs of female *Fmr1* KO mice were tested. The mice were 38-42 days old. Social interaction was assessed in a 30 x 30 x 30 cm chamber, with an overhead camera attached to the lid. To conduct the social interaction test, two mice of the same age, sex, and genotype but from different cages, were both placed into the testing cage. Before the test, neither male nor female mice had been exposed to the opposite sex after weaning. The mice were group-housed and habituated to the behavioral testing room in their respective home cages for 30 minutes before testing. The matched mice were then transferred to the social tracking chamber for testing. Testing sessions were recorded for 10 minutes with Spinview software. The chamber was cleaned with 70% ethanol and left to dry for several minutes before the start of subsequent tests.

**Figure 1.**
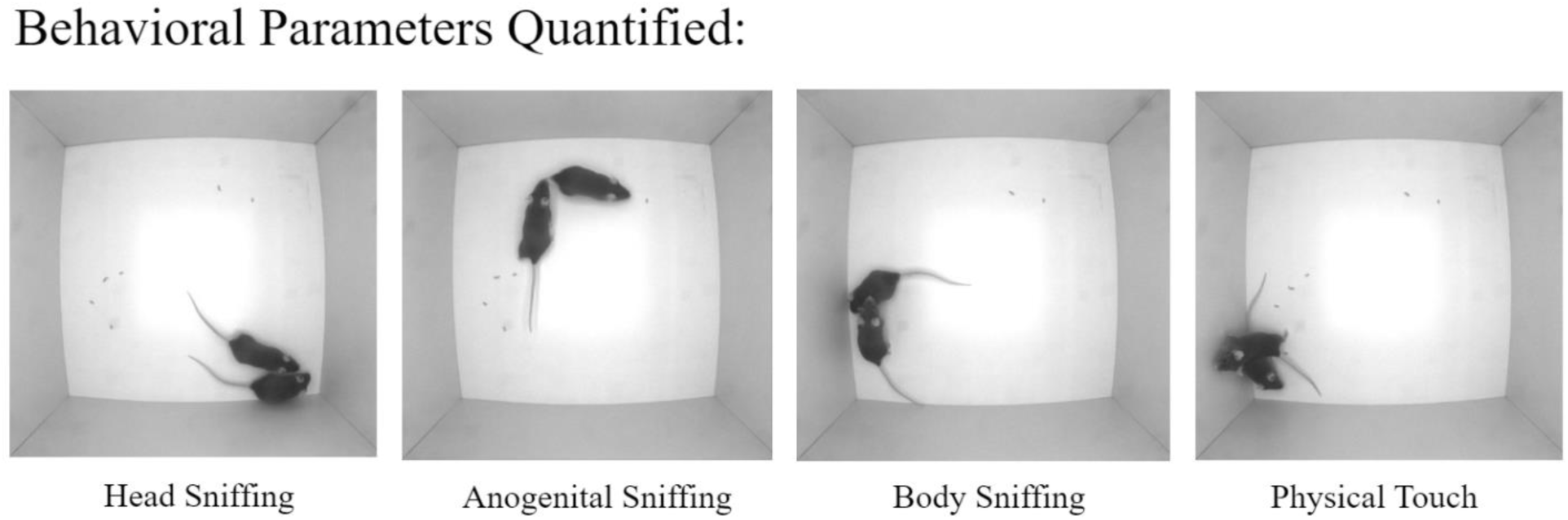
Real time images of behavioral parameters. These four real time images illustrate the parameters quantified, specifically, for head sniffing, anogenital sniffing, body sniffing, and physical touch.

### 2.3 Behavioral Analysis

The coordinates of each mouse’s snout and tail-base, as well as the body mask, were generated using our newly developed marker-less tracking algorithm (Le *et al*., 2024). These parameters were then used to determine social investigation of each mouse towards the other. Specifically, the amount of time the snout of each mouse was directed towards the anogenital, body, or head/face region of the other mouse was quantified. If one mouse’s snout was within a 25-pixel threshold distance from the other mouse’s snout, tail-base, or body, a behavioral label was given (Le *et al*., 2024). This threshold was determined based on the following: head length was measured from the snout to the midpoint of the ears in approximately 200 images; the average head length was determined to be ∼20 mm, corresponding to ∼25 pixels in the analyzed images. Since our approach generates body masks in addition to the snout and tail-base coordinates, we could also determine how much time the pair of mice spent in contact with each other by detecting at what times the masks overlapped. By removing the snout-directed social investigation times from this data, we could determine the amount of time each pair spent touching that was not because of snout-directed investigation. Thus, we analyzed four types of social behaviors including: head sniffing, anogenital sniffing, sniffing of any part of the partner’s body excluding head and anogenital area (body sniffing), and physical touch. For each behavior, we quantified the total time each animal spent in doing that behavior, the occurrence counts of that behavior, the average duration of each occurrence, and the average interval between occurrences. We also used the tracking algorithm to measure the speed as well as distance and trajectory travelled by each animal.

### 2.4 EEG surgery and recording

EEG surgeries and recordings were performed using our established protocol (Cheng & Murari, 2020; Mayengbam *et al*., 2021). For this experiment, 14 female WT mice and 12 female *Fmr1* KO mice were tested. In short, mice were anesthetized with 5% isoflurane and maintained at 1-2% during surgery. Once anesthetized, mice were placed on a heated stereotaxic stage, eyes protected with gel, and the scalp cleaned with 70% ethanol and iodine. Anesthesia depth was confirmed by the absence of toe pinch reflex. The scalp was subsequently removed, connective tissue displaced, and the skull cleaned with hydrogen peroxide. For differential frontal-parietal recordings, three small holes were drilled into the skull at specific coordinates: ground (AP: +2.5 mm, ML: +1.3 mm), left parietal cortex (AP: -2.2 mm, ML: -2.5 mm), and reference electrode in the left frontal cortex (AP: +2.5 mm, ML: -1.3 mm), relative to bregma, above the dura. Electrodes were connected to a miniature connector and secured with glue and dental cement. After surgery, mice received Metacam and post-operative warming until recovery.

EEG recording commenced five days post-surgery to allow for recovery. Prior to recording, the subject mouse underwent brief sedation with isoflurane inhalation before being connected to a standalone EEG device. A 30-minute recording session in the home cage (HC) served as a baseline and was followed by testing in the free social interaction (SI) arena.

### 2.5 EEG data analysis

EEG data analysis followed established methods (Murari *et al*., 2023). Recording quality was manually screened; datasets with obvious electrical noise were excluded. To generate **Fig. 8**, continuous recordings were first segmented into epochs aligned to the onset of the behavioral task. Analysis was performed in MATLAB R2023a (MathWorks) using Welch’s method: signals were segmented into 2-second, 50%-overlapping epochs, and power spectral density (PSD) was computed with a spectral resolution of 0.5 Hz. Band-specific power was calculated by trapezoidal integration of the PSD for delta (1-4 Hz), theta (4-8 Hz), alpha (8-13 Hz), beta (13-30 Hz), and gamma (30-100 Hz), with gamma further subdivided into low (30-60 Hz) and high (60-100 Hz) components. Power values were then normalized to a pre-baseline period (-1 to 0 min relative to task onset) to account for inter-animal variability. Normalized power changes were averaged across animals within each group, and the mean ± SEM values were plotted as shown in **Fig. 8**. Statistical comparisons across time and genotype were performed using two-way repeated-measures ANOVA followed by Tukey’s post hoc tests.

### 2.6 Statistical Analysis

Sample size was determined from previous studies and preliminary data to ensure reliability. Statistical analyses were performed in GraphPad Prism 10.1.0 (GraphPad Software, San Diego, California), with figures generated in Prism and Excel. Analysis details are provided in figure legends and tables. Data are shown as mean ± SEM with α set at 0.05. Two-way ANOVA assessed effects of genotype and sex, followed by Tukey post-hoc tests. For EEG analyses across time in the free social interaction test, a two-way repeated measures ANOVA was used to compare genotype and time, with Tukey post-hoc comparisons.

## 3. Results

Using the marker-less tracking software, we quantified four social behaviors: head sniffing, anogenital sniffing, body sniffing, and physical touch. For each social behavior, we measured the following parameters: the total duration subjects engaged in this behavior, the accumulative time spent in this behavior over the course of the test, the duration of a single event of this behavior, intervals between single events, and count of total events of this behavior. We also quantified the total distances travelled by the subjects as an indication of their locomotor activity levels.

### 3.1 For head sniffing behavior, there were significant sex and genotype differences for all parameters measured

We first quantified head sniffing behavior **(Fig. 2)**. First, we measured the total duration of this behavior **(A)**. There was a main effect of sex observed. The total time spent in this behavior was significantly shorter in females compared to males. There was also a main effect of genotype for this parameter. The accumulative time spent in this behavior followed a gradual increase over time for both sexes and genotypes **(B)**. We then quantified the duration of a single event. There was a main effect of sex **(C)**. Notably, females spent a greater time in a single head sniffing event compared to males of the same genotype. WT females had a longer duration of a single event compared to WT males, and KO females had a longer duration than KO males. A main effect genotype difference was observed, as the WT females spent a greater time in a single event compared to both HET and KO females, and the WT males also had a longer duration compared to KO males. We then quantified the interval between single events **(D)**. We observed a main effect of sex. The interval between single events was longer in females than males of the same genotype. A main effect of genotype was also observed. WT had the longest interval in both sexes, followed by KO. Finally, we measured the count of total events **(E)**. There was a main effect of sex. Females exhibited a lower count than males in the same genotype. Main effect of genotype was also present, as both WT and KO females displayed a lower event count compared to HET females.

**Figure 2.**
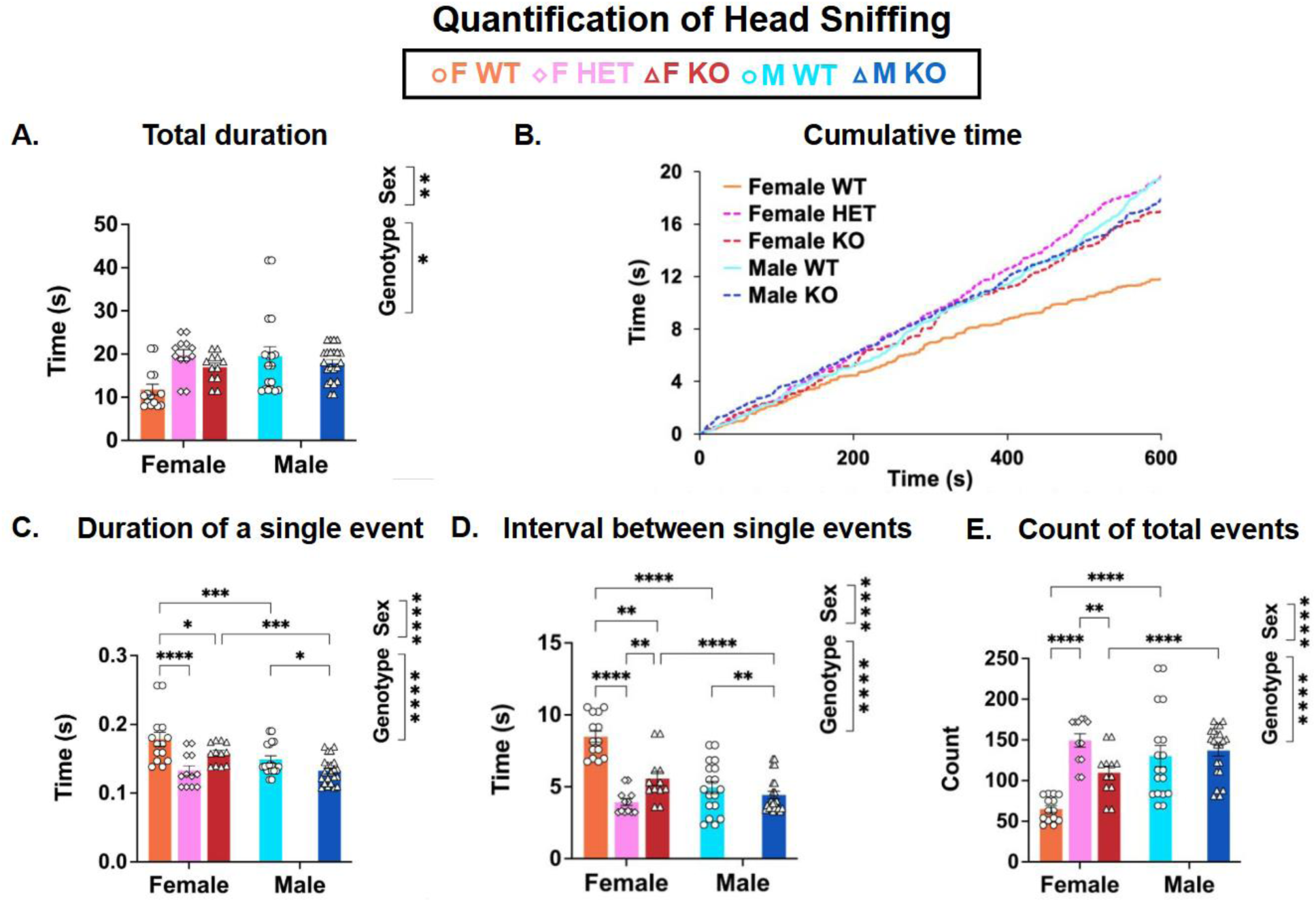
Quantification of head sniffing behavior parameters of juvenile mice in 10-minute recordings. This figure shows the total duration of social interactions (A), cumulative time (B), duration of single event (C), interval between single events (D), and the count of total events in juvenile WT (circles), HET females (diamonds) and *Fmr1* KO (triangles) (E). Juvenile females, WT (n=14), HET (n = 12), and KO (n=12); juvenile males, WT (n=18) and KO (n=22). Data is expressed as mean (bars) ± SEM (error bars) and individual animals (symbols). *p* value was *p* < 0.05.

### 3.2 For anogenital sniffing behavior, there were significant sex differences in all the parameters measured and a genotype difference in all the parameters except duration of a single event

We then quantified anogenital sniffing behavior **(Fig. 3)**. First, we measured the total duration of this behavior **(A)**. There was a main effect of sex observed. There was also a main effect of genotype for this parameter. The duration was shorter in WT females compared to HET females. The accumulative time spent in anogenital sniffing with their partners followed a gradual increase overtime for both sexes and genotypes **(B)**. We then quantified the duration of a single event **(C)**. There was a main effect of sex. Interestingly, the female mice had a longer duration of a single anogenital sniffing event compared to the males. Specifically, both WT and KO females had a longer duration than males of the same genotype. There was no main effect of genotype observed. We then quantified the interval between single events and observed a main effect of sex **(D)**. WT and KO female mice showed longer intervals compared to their male counterparts. Additionally, there was a main effect of genotype. WT females showed a longer interval between single events than both HET and KO females. Further, WT males showed a longer interval than KO males. Finally, we measured the count of total events **(E)**. There was a main effect of sex observed. Both WT and KO females showed a lower number of total events compared to males of the same genotype. A main effect of genotype was also present, as both WT and KO females showed a smaller count than HET females.

**Figure 3.**
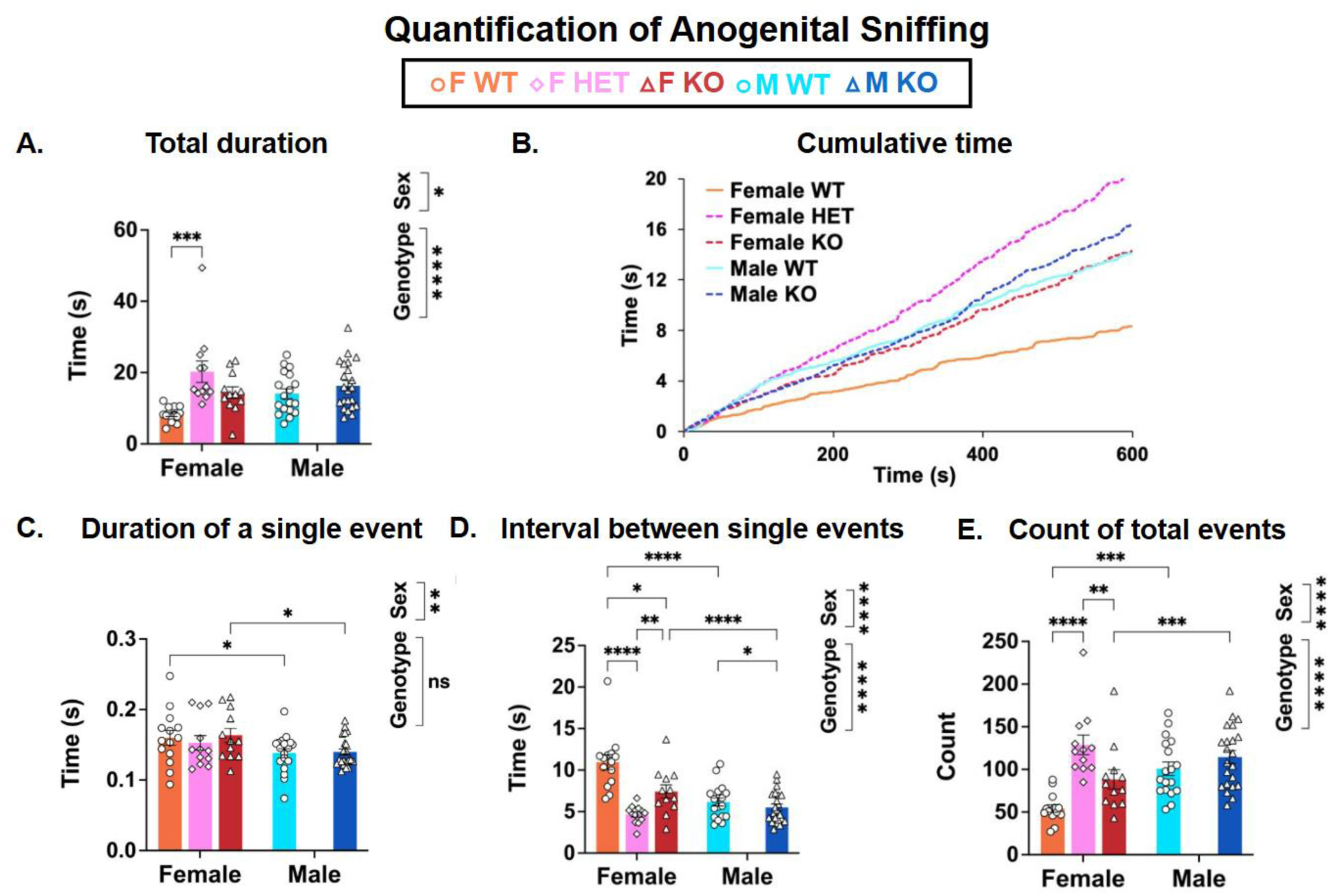
Quantification of anogenital sniffing behavior parameters of juvenile mice in 10-minute recordings. This figure shows the total duration of social interactions (A), cumulative time (B), duration of single event (C), interval between single events (D), and the count of total events in juvenile WT (circles), HET females (diamonds) and *Fmr1* KO (triangles) (E). Juvenile females, WT (n=14), HET (n = 12), and KO (n=12); juvenile males, WT (n=18) and KO (n=22). Data is expressed as mean (bars) ± SEM (error bars) and individual animals (symbols). *p* value was *p* < 0.05.

### 3.3 When quantifying body sniffing behavior, we found a sex difference in each of the parameters except total duration, and further genotype differences in all parameters

We quantified body sniffing behavior (Fig. 4). First, we measured the total duration of this behavior **(A)**. There was no main effect of sex observed. However, there was a main effect of genotype. In general, the total duration spent body sniffing was shorter in WT compared to KO mice. The accumulative time spent followed a gradual increase over time for both sexes and genotypes **(B)**. We then quantified the duration of a single event **(C)**. There was a main effect of sex. There was also a main effect of genotype, as HET females showed a shorter duration of a single event than both WT and KO females. We then quantified the interval between single events **(D)**. There was a main effect of sex observed. Both WT and KO female mice showed longer intervals between single body sniffing events compared to their respective male counterparts. There was also a notable main effect of genotype. Both WT and KO females showed a shorter interval between events compared to the HET females. WT males showed a greater interval compared to KO males. Finally, we measured the count of total events **(E)**. There was a main effect of sex. Females showed a smaller number of total events compared to males of the same genotype. There was also a main effect of genotype. WT females exhibited a significantly smaller count compared to both HET and KO females. WT males also exhibited a smaller count than KO males.

**Figure 4.**
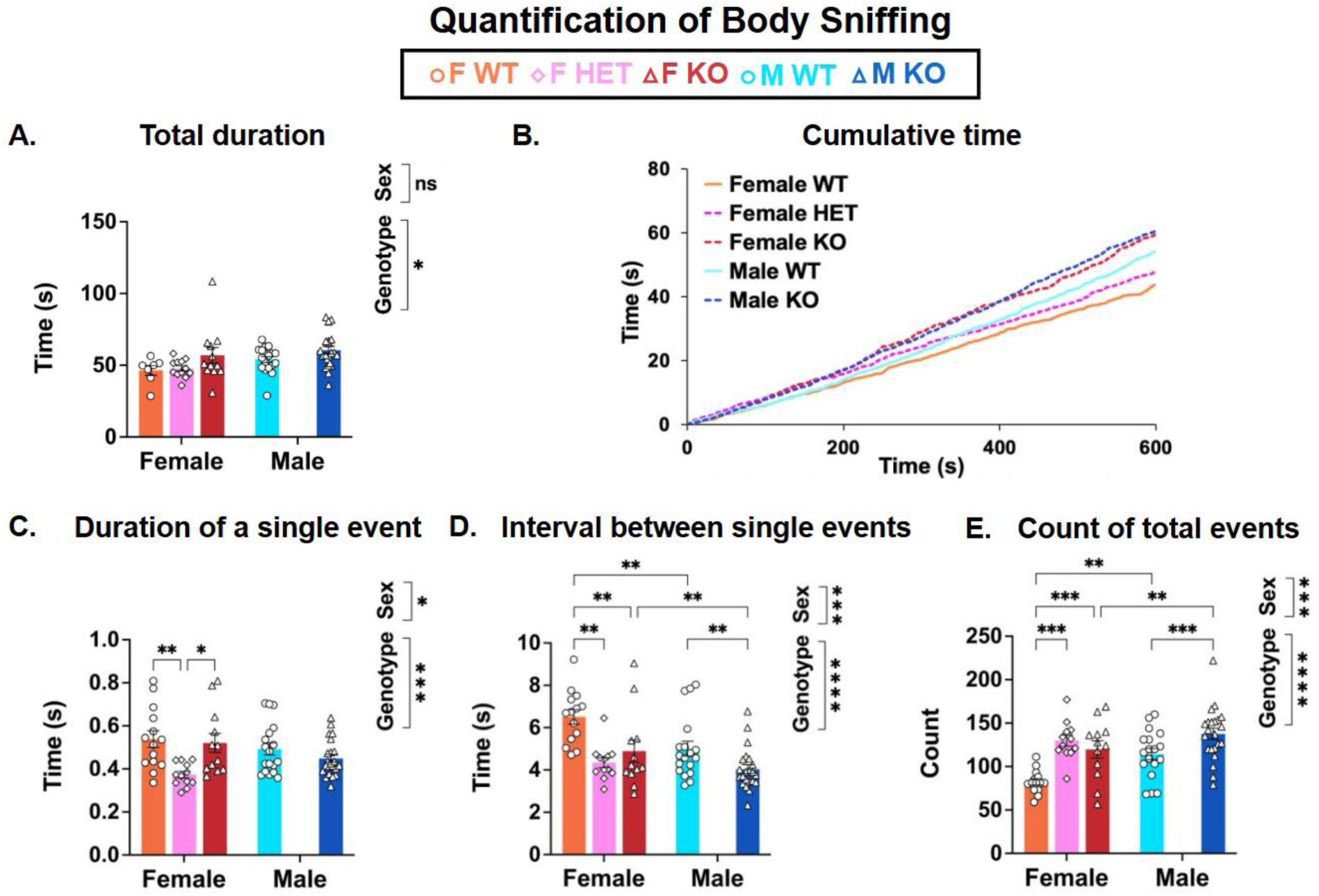
Quantification of body sniffing behavior parameters of juvenile mice in 10-minute recordings. This figure shows the total duration of social interactions (A), cumulative time (B), duration of single event (C), interval between single events (D), and the count of total events in juvenile WT (circles), HET females (diamonds) and *Fmr1* KO (triangles) (E). Juvenile females, WT (n=14), HET (n = 12), and KO (n=12); juvenile males, WT (n=18) and KO (n=22). Data is expressed as mean (bars) ± SEM (error bars) and individual animals (symbols). *p* value was *p* < 0.05.

### 3.4 During physical touch behavior, there were sex and genotype differences observed for each of the parameters

We quantified physical touch behavior (Fig. 5). First, we measured the total duration of this behavior **(A)**. There was a main effect of sex observed. Specifically, both WT and KO females showing shorter durations compared to males of the same genotype. Further, there was a main effect of genotype. WT females showed less total interaction than both HET and KO females. WT males also showed fewer total interactions than KO males. The accumulative time spent in physical touch with their partners followed a gradual increase over time for both sexes and genotypes **(B)**. We then quantified the duration of a single event **(C)**. There was a main effect of sex, as females showed slightly longer duration of single physical touch event compared to males. There was also a main effect of genotype. WT females showed significantly more time in a single event compared to HET females. We also quantified the interval between single events **(D)**. There was a main effect of sex. Both WT and KO female mice showed longer interval between single events compared to their respective male counterparts. There was also a main effect of genotype. WT females showed significantly more time between single events compared to HET and KO females. WT males showed a longer interval than KO males. Finally, we measured the count of total events **(E)**. There was a main effect of sex. Both WT and KO females showed a lower number of total events compared to males of the same genotype. There was also a main effect of genotype. In females, both WT and KO mice showed a smaller number of total events than HET mice. Further, WT males displayed less events than KO males.

**Figure 5.**
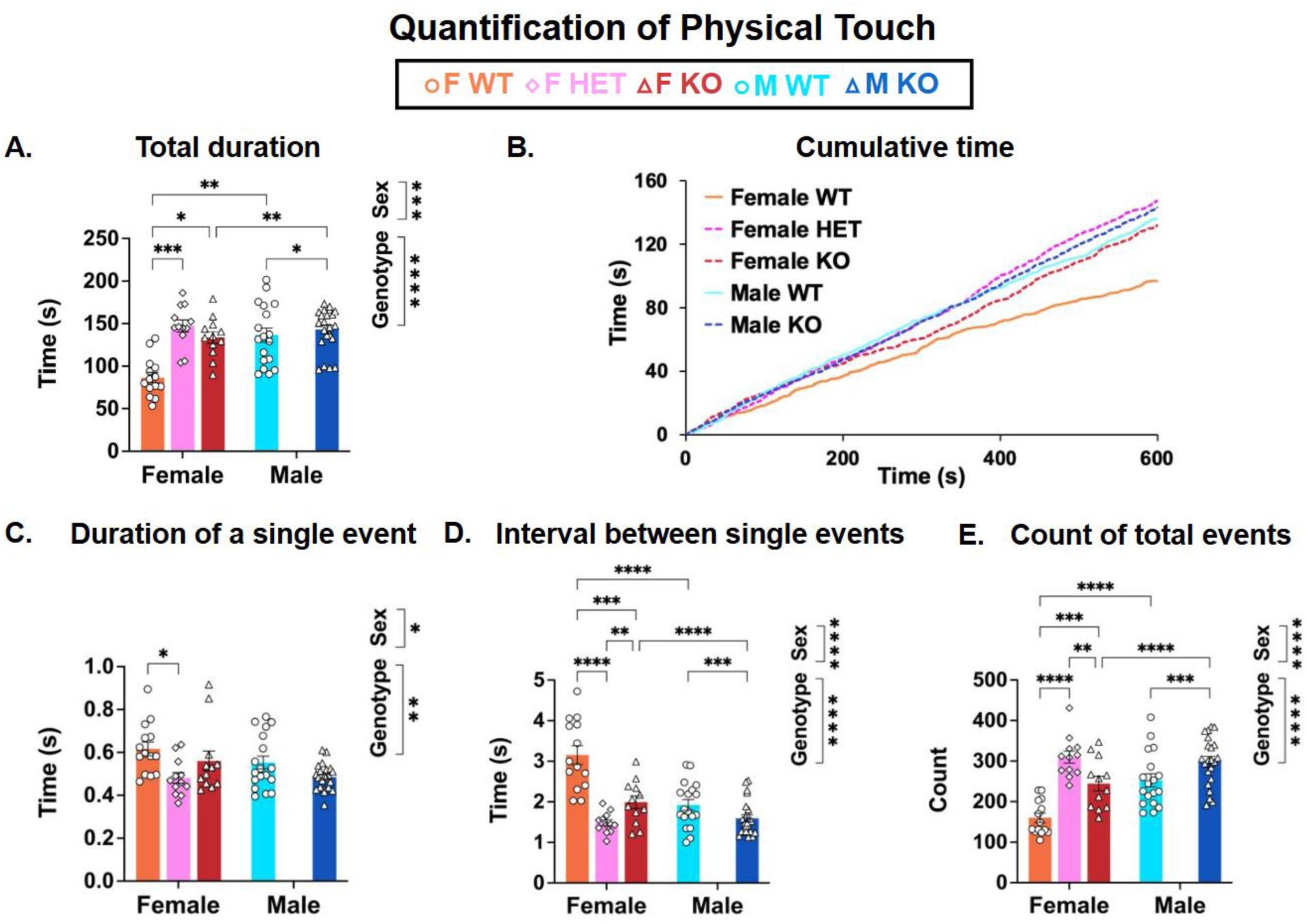
Quantification of physical touch behavior parameters of juvenile mice in 10-minute recordings. This figure shows the total duration of social interactions (A), cumulative time (B), duration of single event (C), interval between single events (D), and the count of total events in juvenile WT (circles), HET females (diamonds) and *Fmr1* KO (triangles) (E). Juvenile females, WT (n=14), HET (n = 12), and KO (n=12); juvenile males, WT (n=18) and KO (n=22). Data is expressed as mean (bars) ± SEM (error bars) and individual animals (symbols). *p* value was *p* < 0.05.

### 3.5 When considering the distance travelled by the mice, there were sex and genotype differences observed

The distance traveled was then quantified (Fig. 6). First, we measured the total duration of this behavior **(A)**. There was a main effect of sex observed. Specifically, WT and KO females travelled more than their respective male counterparts. There was also a main effect of genotype observed. Both WT and KO females travelled less than HET females.

**Figure 6.**
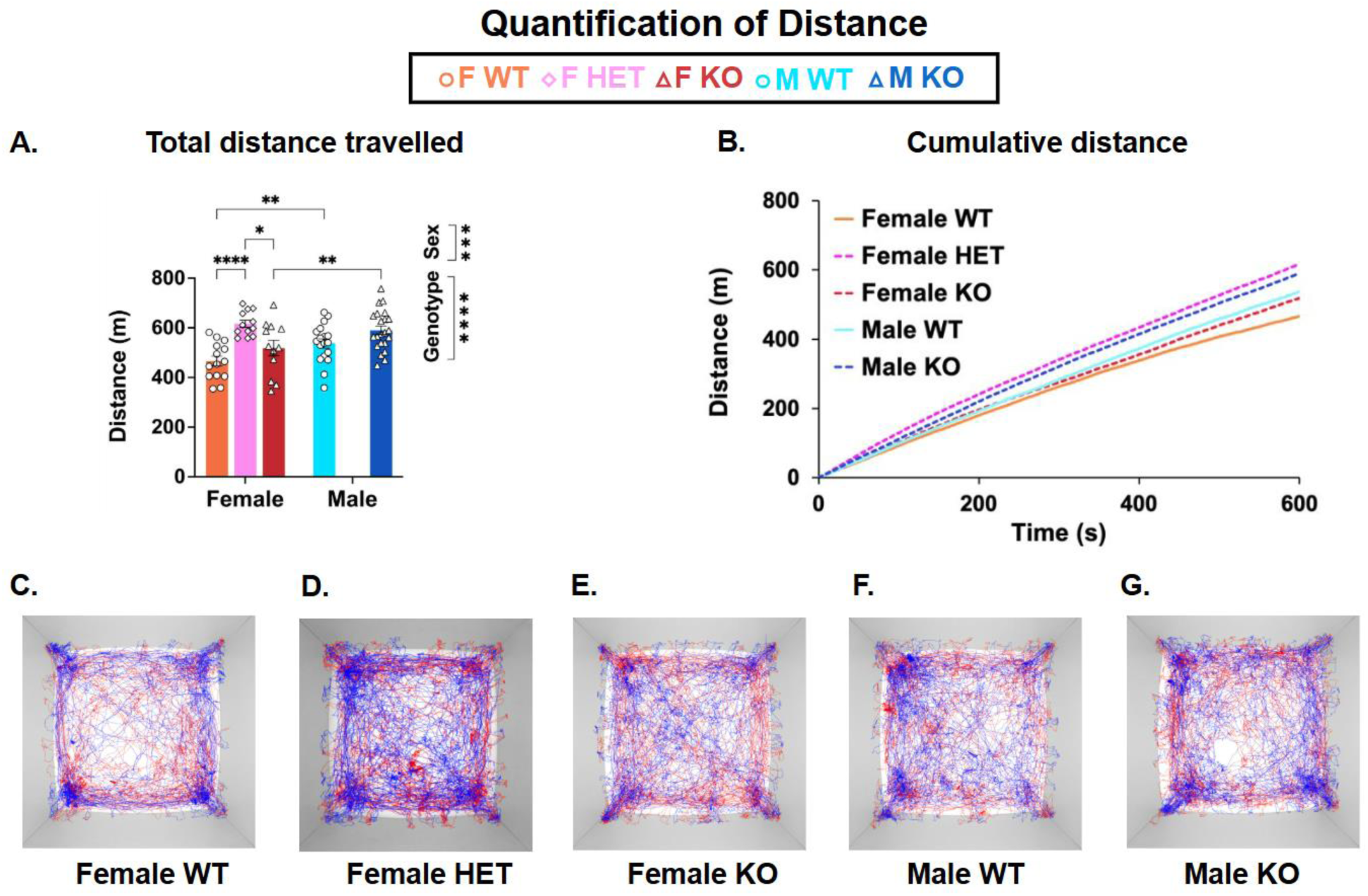
Quantification of distance travelled by juvenile mice in 10-minute recordings. This figure shows the total distance travelled (A), cumulative distance (B), and tracking trajectories (C-G). Juvenile females, WT (n=14), HET (n = 12), and KO (n=12); juvenile males, WT (n=18) and KO (n=22). Data is expressed as mean (bars) ± SEM (error bars) and individual animals (symbols). *p* value was *p* < 0.05.

The accumulative distance travelled followed a gradual increase over time for both sexes and genotypes **(B)**. Tracking trajectories for each group were quantified by the algorithm software, with blue lines representing the body center of one mouse and red tracing the other mouse **(C-G)**.

**Table I.** Data summary of social behavioral parameters.

|  |  | Sex |  | Genotype |  |  |  |
| --- | --- | --- | --- | --- | --- | --- | --- |
|  |  | WT<br>females<br>vs. WT<br>males | KO<br>females<br>vs. KO<br>males | WT<br>females<br>vs. HET<br>females | WT<br>females<br>vs. KO<br>females | HET<br>females<br>vs. KO<br>females | WT<br>males<br>vs. KO<br>males |
| **Total**<br>**duration** | Head<br>sniffing |  |  |  |  |  |  |
|  | Anogenital<br>sniffing |  |  | *** |  |  |  |
|  | Body<br>sniffing |  |  |  |  |  |  |
|  | Physical<br>Touch | ** | ** | *** | * |  | * |
| **Duration**<br>**of single**<br>**event** | Head<br>sniffing | *** | *** | **** | * |  | * |
|  | Anogenital<br>sniffing | * | * |  |  |  |  |
|  | Body<br>sniffing |  |  | ** |  | * |  |
|  | Physical<br>Touch |  |  | * |  |  |  |
| **Interval**<br>**between**<br>**single**<br>**events** | Head<br>sniffing | **** | **** | **** | ** | ** | ** |
|  | Anogenital<br>sniffing | **** | **** | **** | * | ** | * |
|  | Body sniffing | ** | ** | ** | ** |  | ** |
|  | Physical Touch | **** | **** | **** | *** | ** | *** |
| Count of total events | Head sniffing | **** | **** | **** |  | ** |  |
|  | Anogenital sniffing | *** | *** | **** |  | ** |  |
|  | Body sniffing | ** | ** | *** | *** |  | *** |
|  | Physical Touch | **** | **** | **** | *** | ** | *** |

### 3.6 Time-resolved neural oscillatory dynamics in WT and *Fmr1* KO mice during social interaction

Prior to analyzing time-resolved changes in EEG dynamics during social interaction, we first confirmed data quality and baseline neural activity. As illustrated in Fig. 7, representative raw EEG traces and power spectra from WT and *Fmr1* KO female mice during the resting condition demonstrated the stability and reliability of our recordings.

**Figure 7.**
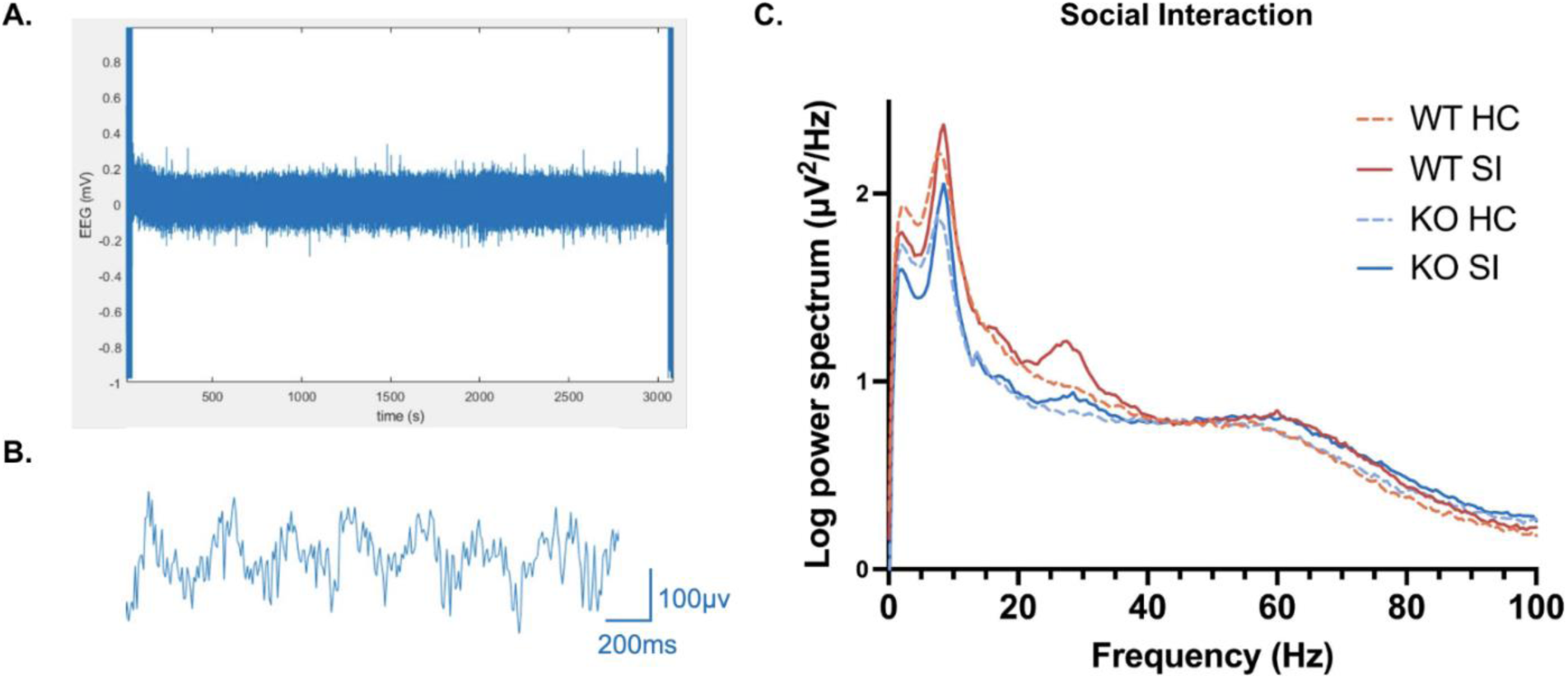
Power spectra of KO and WT of female juvenile mice in the social interaction at each condition. EEG raw signal from 50 minutes of recording. The noise in the first and last 120 seconds (about 2 minutes) shows the time when the mice were anesthetized, and the EEG recording device is connected to the mouse head cap (A). Example of EEG signal (B). Power spectra of KO (n=12) and WT (n=14) female mice in the home cage (HC) and social interaction (SI) arena (C).

To assess how neural oscillations evolved during social interaction, we analyzed normalized EEG power at discrete time points relative to the onset of interaction (-1, +1, +3, +5, +7, and +9 min; Fig. 8). Power values were expressed relative to each group’s average at baseline (-1 min), thereby standardizing the baseline to 1. This normalization minimized inter-animal variability and allowed us to focus on task-induced changes.

**Figure 8.**
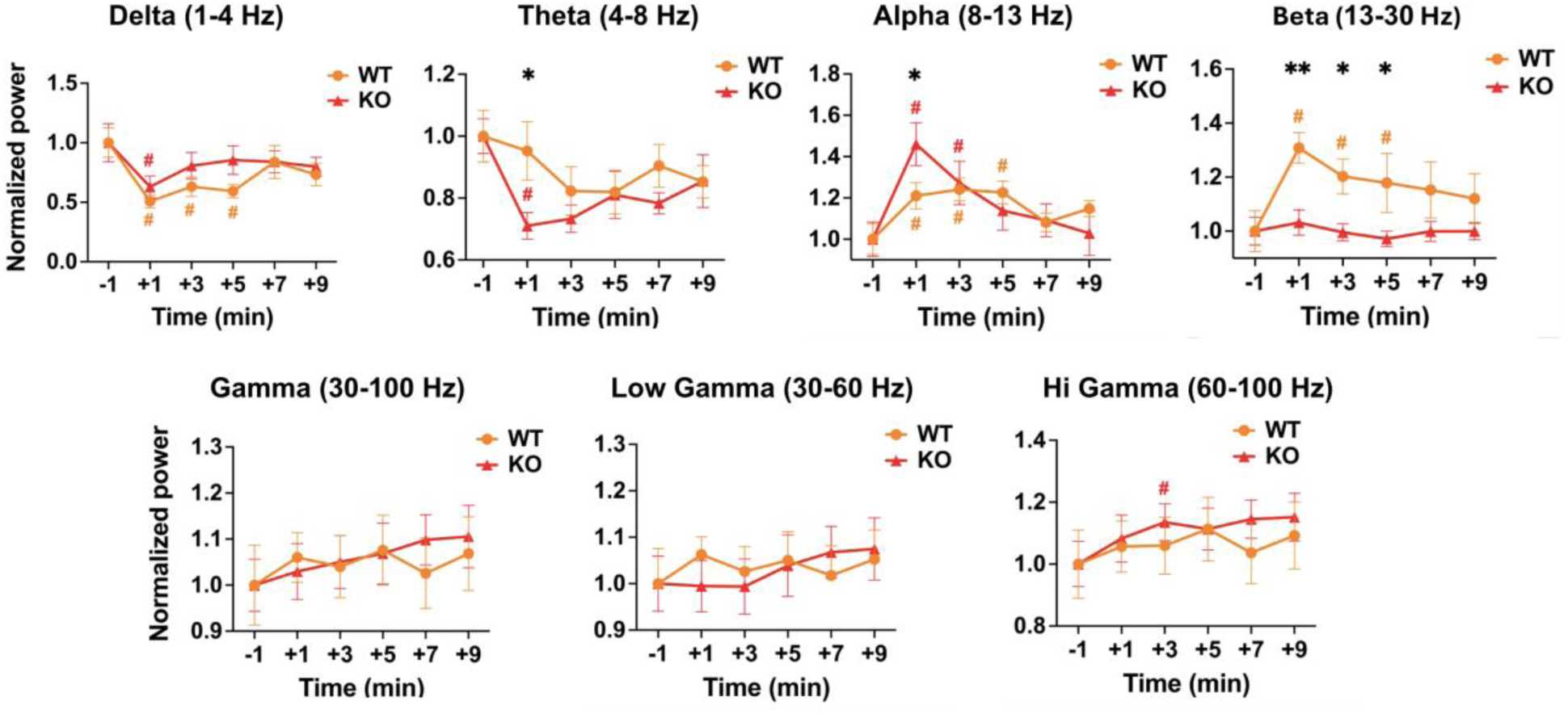
Time-point analysis of normalized relative EEG power across frequency bands in WT and KO mice during social interaction tests. Plots show the mean ± SEM relative power at each indicated time point relative to the onset of social interaction (-1, +1, +3, +5, +7, +9 min). Relative power values were used to normalize EEG data within each frequency band. Statistical analysis was performed using two-way repeated measures ANOVA, followed by Tukey’s multiple comparison test. Genotype differences at specific time points are indicated by asterisks (*: p < 0.05, **: p < 0.01). Time point differences (relative to -1 min) are marked by section sign (#): orange # for WT, and red # for KO.

Two-way repeated-measures ANOVA revealed significant main effects of genotype and time, particularly in the lower frequency ranges. In the theta band, KO mice exhibited reduced normalized power compared to WT at +1 min. In contrast, KO mice showed elevated alpha power relative to WT during the same early phase. The most robust effect was observed in the beta band, where KO mice displayed consistently lower normalized power across multiple post-onset intervals (+1, +3, +5 min).

By comparison, no significant genotype differences emerged in the delta or gamma ranges, and both groups exhibited relatively stable normalized power in higher frequencies throughout the test session. Together, these results indicate that *Fmr1* KO mice show frequency-specific alterations in oscillatory dynamics during social interaction, with the strongest effects occurring in theta, alpha, and beta bands during the initial minutes of engagement.

### 3.7 Social behaviors did not completely correlate with locomotor activity levels but reflected genotype- and sex-specific traits

To examine whether locomotor activity influenced social behaviors, we conducted correlation analyses between the total distance traveled and selected behavioral parameters. Specifically, we assessed correlations with (1) the count of total events (**Table II**) and (2) the duration of a single event (**Table III**) for each social behavior. All p-values were adjusted for multiple comparisons across the entire table using the Benjamini-Hochberg false discovery rate procedure, and only correlations that remained significant after correction are reported below.

**Table II.**
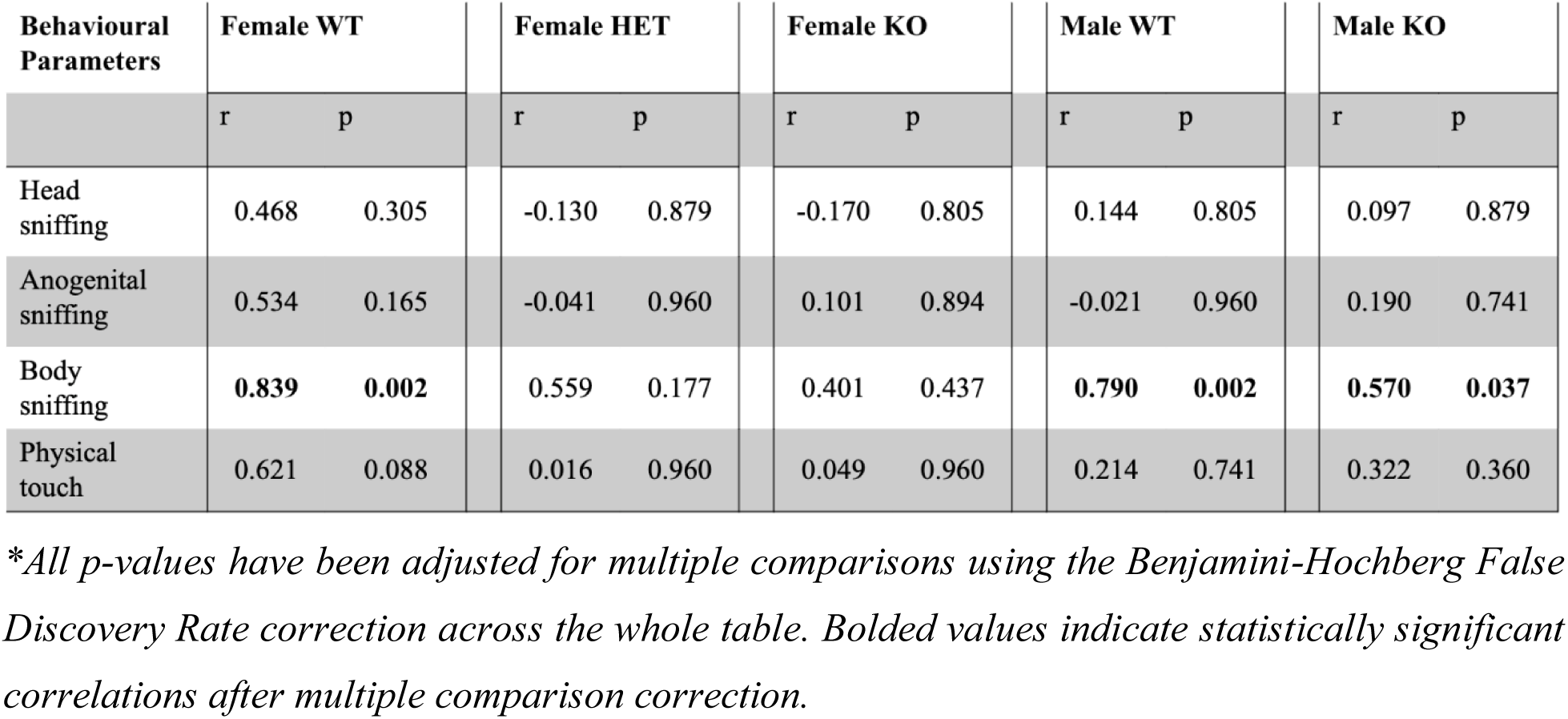
Pearson correlation (r) and p values between the total distance travelled and the count of total events of each social behavior.

**Table III.** Pearson correlation (r) and p values between the total distance travelled and the duration of a single event for each social behavior.

| Behavioural Parameters | Female WT |  | Female HET |  | Female KO |  | Male WT |  | Male KO |  |
| --- | --- | --- | --- | --- | --- | --- | --- | --- | --- | --- |
|  | r | p | r | p | r | p | r | p | r | p |
| Head sniffing | -0.435 | 0.178 | -0.658 | 0.057 | 0.337 | 0.356 | 0.042 | 0.896 | -0.337 | 0.178 |
| Anogenital sniffing | **-0.753** | **0.008** | -0.221 | 0.057 | -0.352 | 0.349 | 0.087 | 0.812 | 0.030 | 0.896 |
| Body sniffing | -0.544 | 0.099 | -0.577 | 0.099 | **-0.875** | **0.001** | **-0.797** | **0.001** | **-0.724** | **0.001** |
| Physical touch | -0.533 | 0.099 | -0.532 | 0.125 | **-0.759** | **0.014** | **-0.780** | **0.001** | -0.389 | 0.125 |

*\*All p-values have been adjusted for multiple comparisons using the Benjamini-Hochberg False Discovery Rate correction across the whole table. Bolded values indicate statistically significant correlations after multiple comparison correction.*

Overall, locomotor activity was only modestly related to social behaviors. For event counts (**Table II**), no significant correlations were found for head, anogenital sniffing or physical touch. By contrast, body sniffing showed a robust positive correlation with distance traveled in both male WT (r= 0.790, p = 0.002) and female WT mice (r=0.839, p=0.002), and a weaker but still significant correlation in male KO mice (r=0.570, p=0.037). For physical touch, a significant positive correlation was detected only in female WT mice.

For event duration (**Table III**), correlations tended to be negative or nonsignificant. In body sniffing, total distance traveled was negatively correlated with the duration of individual events in male WT (r=-0.797, p=0.001), male KO (r=-0.724, p=0.001), and female KO mice (r=-0.875, p=0.001). Similarly, a negative correlation was observed for physical touch in male WT (r=-0.780, p=0.001) and female KO mice (r=-0.759, p=0.014). These patterns suggest that more mobile animals engaged in briefer but more frequent interactions. Taken together, these analyses indicate that locomotor hyperactivity contributed partially to variation in certain social behaviors - particularly body sniffing - but did not account for the overall genotype- and sex-specific behavioral patterns observed.

## 4. Discussion

### 4.1 Social interaction in the *Fmr1* KO mouse model of FXS

Social interactions are a foundational anchor in society and serve as the building block to create a plethora of stable networks and relationships with others. It encompasses multiple individuals exhibiting a diverse and complex range of behaviors affected by their individual emotional states and motivations (Jabarin *et al*., 2022). Differences in social behaviors, especially social anxiety and withdrawal, are prominent features of FXS. In addition, the underlying mechanisms behind the differences in social behavior observed in FXS patients remains unclear, hence there are no effective interventions that can help patients overcome social challenges. As a result, having a precise, comprehensive characterization of social behavior is necessary to understand the neurobiological basis that underlies its complexities.

Several studies have shown that *Fmr1* KO mice can be used to replicate some of the behavioral defects of autism and FXS (Moy *et al*., 2009; Liu & Smith, 2009). Therefore, in this study we compared the social behaviors of both female and male mice of the *Fmr1* KO mouse model of FXS with HET and WT mice through the analysis of several behavioral parameters at the juvenile stage. Juvenile mice provide a unique opportunity to improve our understanding of the importance of early intervention and mitigation of long-term impacts, since this age is considered a sensitive period of the brain development and behavioral plasticity in both mice and humans (Petroni *et al*., 2022; Liu & Smith, 2009). Moreover, studies in humans have shown that young children with FXS may provide a unique understanding of the early signs of developing multiple psychiatric disorders, such as ASD, social anxiety, and generalized anxiety disorder as they grow older (Black *et al*., 2021).

In mice, socio-emotional interactions are mediated by various modalities such as visual, olfactory, and auditory senses (Jabarin *et al*., 2022). Sniffing is a key active behavior to acquire odors and pheromones exhibited during social behaviors (Wesson, 2013). The mechanism by which mice communicate is not fully understood, however, exploration of a novel context through sniffing is a measurement for the arousal state of the animal and is relevant to information exchange between brain areas (Jabarin *et al*., 2022). In mice, sniffing behavior and close physical contact appear to contribute to communication of information and social hierarchy (Silverman *et al*., 2010). Although some sniffing behavior, like anogenital sniffing, has been correlated with the collection of odor information and sexual interest, the reasons underlying other individual sniffing behaviors remain unexplored (D’Amato & Pavone, 1992).

Sociability studies, like the three-chambered sociability task, were used in previous studies to study the behavior of a subject mice when allowed to interact with a novel mouse against a novel object (Yang *et al*., 2011). In general, studies that used the three-chambered sociability tests to study the behavior of *Fmr1* KO mice observed varying social behaviors (Kazdoba *et al*., 2014). Some studies observed that *Fmr1* KO mice were similar to their WT counterparts and preferred to interact with the novel mouse rather than the novel object (Kazdoba *et al*., 2014; Liu & Smith, 2009; Pietropaolo *et al*., 2011; Liu *et al*., 2011). Other studies observed that *Fmr1* KO mice displayed increased social interaction (Spencer *et al*., 2005). One study noted enhanced locomotor activity and increased social interest in the three-chambered sociability task in *Fmr1* KO mice which may be explained by the mice’s more frequent interaction with the stranger mouse relative to the WT mice (Sørensen *et al*., 2015). On the other hand, other studies concluded that *Fmr1* KO mice may not have any preference for the novel mouse over the novel object, which is also seen with decreased duration of sniffing events of the novel mouse in the three chambered sociability tests (Pietropaolo *et al*., 2011). Another study mentions that it is unlikely that hyperactivity reflects the differences in results during the 3-chamber test because there were no differences in crossing the arenas during the habituation and testing phases (Wesson, 2013). Rather, a more likely explanation is that the differences in social interaction are because of greater arousal in the *Fmr1* KO mice than for the WT, making it difficult for them to stay in contact with the stimulus mouse (Wesson, 2013).

Deep learning approaches, as displayed by the software used in the present study, enabled the detection and tracking of changes in the positions of defined body parts, that allowed for the investigation of social interactions in a more natural setting (Kuo *et al*., 2022). In this study using the FXS mouse model, the length of social interaction following initiation of an approach depends on the willingness of both mice, as discrepancies in the cooperation or interest of a mouse will result in a shorter interaction. Our results on male and female *Fmr1* KO, HET female, and WT mice on a C57BL/6J background were complex, in terms of the level of social interactions in a 1-to-1 setting. Based on the analysis of all four behaviors (head sniffing, anogenital sniffing, body sniffing, and physical touch), there were robust sex differences: females exhibited a longer duration in a single interaction event as well as longer intervals between events; in contrast, the male mice had significantly more interaction counts than the female cohorts.

In terms of genotype differences, in all the social behaviors we quantified, the KO mice displayed significantly more interaction events than the WT mice, especially comparing WT and KO females, however HET females displayed the highest count. In a study by McNaughton *et al*. (2008), where adult male mice of age 8-10 months were tested in the three-chambered apparatus, *Fmr1* KO mice had shorter average duration of nose contact interactions and higher levels of grooming during initial interactions. These findings are consistent with the findings in the present study, where KO mice displayed a shorter duration of a single event during head sniffing, compared to the WT counterparts, and may reflect a common feature in this mouse model of FXS. In the study by Petroni *et al*. (2022), showed that HET females displayed a hyper-social phenotype at 7-8 weeks of age, characterized by increased affiliative behaviors during a 3-minute direct social interaction test (Petroni *et al*., 2022). Affiliative behaviors included sniffing the partner’s head, snout, anogenital region, or other body parts; physical contact such as crawling over or under the partner’s body; and allogrooming (Petroni *et al*., 2022). Nonsocial activities were also recorded, such as rearing, digging, and self-grooming (Petroni *et al*., 2022). Initially, we predicted that HET mice would exhibit an intermediate behavior between WT and KO mice, due to the partial gene expression in HET mice. However, the HET mice unexpectedly displayed social behaviors similar to or exceeding those of KO mice, when compared with WT mice. The results suggest that gene dosage of *Fmr1* contributes to social behavior phenotype we measured here in a complex way, which may be related to the mosaic expression of *Fmr1* in the HET females and downstream compensatory mechanisms. Almost all female FXS patients are heterozygous. Our observations thus raise the question of whether gene dosage of *Fmr1* alone could explain the specific traits of female FXS patients that are often described as being milder than the male patients.

### 4.2 Oscillatory Disruptions Reflecting Circuit-Level Dysfunction in *Fmr1* KO Females

Our EEG recordings during social interaction revealed distinct frequency-specific perturbations in female *Fmr1* KO mice. Specifically, they showed reduced theta power, elevated alpha power, and pronounced suppression of beta power in the initial minutes of social engagement, while delta and gamma bands remained unaltered. This profile suggests that oscillatory alterations in *Fmr1* KO mice are selective, but not global. Reduced alpha and increased theta power have been reported in clinical EEG studies of individuals with FXS. Notably, these changes were observed during resting-state recordings (Van der Molen & Van der Molen, 2013). In contrast, our data were obtained while mice were actively engaged in free social interactions - a condition more akin to task-evoked brain activity. Further studies to expand EEG recording in different settings would likely provide additional insight into neural network mechanisms in FXS.

On the other hand, changes in gamma power have been consistently observed in both human FXS patients and rodent models, typically manifesting as pronounced increases at the resting stage (Wilkinson & Nelson, 2021; Wang *et al*., 2017). However, our data indicate that in female KO mice, theta-beta band dynamics are especially sensitive to social context. Functionally, beta oscillations are critical for long-range cortical communication, social cognition, and top-down control, which are all processes disrupted in FXS (Engel & Fries, 2010; Kilavik *et al*., 2013). The sustained beta suppression observed in KO mice likely reflects changes in these integrative networks during periods of social salience. Similarly, fluctuations in theta and alpha bands, which typically mark attentional and stimulus-processing engagement, point to differences in the processing of social cues and attention (Cavanagh & Frank, 2014).

### 4.3 Contribution of Locomotion to Social Interactions

Our correlation analyses suggest that locomotor hyperactivity in FXS mice may contribute to, but does not fully explain, the increased social interaction events observed in this study. Increased interaction events likely involve factors beyond motility. Of the four parameters analyzed - head sniffing, anogenital sniffing, body sniffing, and physical touch - body sniffing showed the strongest positive correlations with locomotor activity across genotypes and sexes, whereas head sniffing and anogenital sniffing displayed minimal correlations with total distance travelled. These findings indicate that differences in social behavior are not solely attributable to variations in locomotor activity but reflect genotype- and sex-specific traits.

### 4.4 Limitations and Future Directions

A challenge in behavioral analysis pertains to group dynamics. This study focused on interactions between two mice, so the behavior of mice in this dyadic circumstance is not accurately predictable. This is because certain aspects of social behavior are influenced by interactions that do not just involve another individual. The bulk of research on animal models relies on simple behavioral paradigms, thus hindering our understanding of the neural mechanisms governing other aspects of social behavior in more complex settings. For example, emotions greatly influence behavior, yet most studies, including this one, do not include a reliable means for assessing the emotional state of the mice undergoing the social interaction test. Further, the social behavior of a tested mouse may be affected by variables including established dominance, social rank, and individual temperaments (Mineur *et al*., 2006). Thus, performance in these tests may be influenced by multiple variables.

Particularly, we have yet to understand the contribution of social motivation versus anxiety to the observed differences between WT and KO mice. This is an important issue because social motivation and anxiety co-exist in FXS. In studies of FXS patients, Hong *et al*. (2019) used emotional face gaze analysis in patients between 5-30 years of age to study social interest and anxiety symptomology and found that both occur and contribute to social difficulties in individuals with FXS. The study found that individuals with FXS displayed increased gaze aversion but did not show reduced social interest (Hong *et al*., 2019). Interestingly, it also found that greater social preference was correlated with more severe anxiety in FXS patients (Hong *et al*., 2019). This may be due to socially anxious individuals being more hypervigilant to surrounding stimuli or because individuals with FXS that have a higher level of social interest are more susceptible to social anxiety compared to their less socially cognizant peers (Hong *et al*., 2019). These findings are overall consistent with studies reporting that individuals with FXS show behaviors suggestive of a willingness and motivation to interact with others, despite their social communication impairments and anxiety (Crawford *et al*., 2017).

Additionally, a key limitation was the use of broad surface EEG recordings with limited spatial specificity; future studies employing source-localized or laminar techniques could clarify which neural circuits are most affected. It will also be crucial to determine whether these oscillatory patterns change across development or differ by sex, given emerging evidence for sex-dependent EEG phenotypes in FXS.

Future research could benefit from exploring the role of autonomic nervous system dysregulation in FXS-related social avoidance behaviors. Several researchers have suggested that social avoidance behavior in FXS may result from a dysregulation of the sympathetic and parasympathetic nervous systems, providing a new explanation for why people with FXS often seem overly anxious or agitated during social interactions (Hall *et al*., 2009). Another study by Belser and Sudhalter (1995) reported that tonic skin conductance, which reflects activity of sweat glands, of male subjects with FXS were significantly elevated during direct eye contact with others, indicating potential maladaptive responses to social cues. Additionally, a study by Miller *et al*. (1999) found that electrodermal responses to sensory stimuli tests, also indicating sweat gland activity and further sympathetic system activity, were elevated. Further, inversely proportional ratios to the levels of FMRP were detected, such that people with higher levels of FMRP were associated with a more typical electrodermal response (Miller *et al*., 1999). Several studies have observed heightened physiological arousal in FXS patients during cognitively demanding tasks (Roberts *et al*., 2001; Keysor *et al*., 2002). These observations include significantly elevated heart rates, indicative of sympathetic nervous system activity, and significantly lower vagal tone, a measure of parasympathetic nervous system activity (Roberts *et al*., 2001; Keysor *et al*., 2002). These differences are consistently seen in individuals with FXS when compared to a group of age-matched typically developing children, both at baseline and during completion of cognitive tasks (Roberts *et al*., 2001; Keysor *et al*., 2002). Based on these studies, it is likely that investigating arousal states of animal models can provide new understanding of social behaviors in both typical and altered states.

## 5. Conclusions

This study demonstrated the differences associated with various social behaviors in the juvenile *Fmr1* KO mouse model of FXS. Together, our data provides evidence for the presence of behavioral discrepancies in female mice, highlighting the importance of considering age and sex differences when using this mouse model. These results also suggest that investigating the early post-natal phases is crucial for potential preventative interventions. Future studies are necessary to reveal the cognitive and neural processes governing normal social behaviors and can aid in understanding how these mechanisms may produce differences in social behavior.

## 8. Data availability

All data supporting the findings of this study are fully presented within the manuscript. Additionally, the raw data generated and analyzed during the current study are available from the primary and the corresponding author upon reasonable request.

## List of Abbreviations Used

FXS: Fragile X Syndrome
*Fmr1*: Fragile X Messenger Ribonucleoprotein 1
FMRP: Fragile X Messenger Ribonucleoprotein
ADHD: Attention deficit hyperactivity disorder
ASD: Autism spectrum disorder
KO: Knockout
WT: Wildtype
EEG: Electroencephalography
HET: Heterozygous
OSERR: Open-Source Electrophysiology Recording system for Rodents
HC: Home cage
SI: Social interaction
PSD: Power spectral density

